# Multimodal approach to characterize surgically removed epileptogenic zone from patients with focal drug-resistant epilepsy: from operating room to wet lab

**DOI:** 10.1101/2025.04.15.648977

**Authors:** Jenni Kyyriäinen, Adriana Della Pietra, Mastaneh Torkamani-Azar, Mireia Gómez-Budia, Polina Abushik, Nataliia Novosolova, Henri Eronen, Omar Narvaez, Ekaterina Paasonen, Vera Lezhneva, Anssi Pelkonen, Liudmila Saveleva, Antti Huotarinen, Ilya Belevich, Eija Jokitalo, Kuopio Epilepsy Center Epilepsy Surgery Group, Leena Jutila, Tuomas Rauramaa, Arto Immonen, Ville Leinonen, Jussi Tohka, Olli Gröhn, Reetta Kälviäinen, Tarja Malm, Alejandra Sierra

## Abstract

**Objective:** We have established a comprehensive sample handling protocol designed for the multiscale assessment of epileptogenic tissue. This protocol aims to identify novel therapeutic targets and enhance the diagnosis and stratification of patients with drug-resistant epilepsy, thereby optimizing their treatment with anti-seizure medications and surgical interventions.

**Methods:** Patients with drug-resistant focal epilepsy, recommended for surgical treatment, are recruited after detailed multidisciplinary preoperative evaluation at the Epilepsy Center at Kuopio University Hospital in Finland. A day before the resective surgery, patients undergo magnetic resonance imaging (MRI) including advanced methodologies. During the surgery, each piece of resected tissue is placed under oxygenation on ice-cold artificial cerebral spinal fluid-solution. The pieces are then immediately transported to the laboratory, assessed by a neuropathologist, and sliced for both clinical diagnosis and research. Two adjacent slices are provided for research and are sent to the University of Eastern Finland.

**Results:** The developed sample handling protocol provides the opportunity for detailed characterization of the tissue from the same patient using emerging imaging, electrophysiology, and molecular biology technologies. We have optimized the conditions for preserving the resected tissue alive for electrophysiological measurements and simultaneously making possible *ex vivo* studies including multi-omics acquisition, electron microscopy, histology, and MRI. Our protocol enables the mapping of functional readouts to structural and molecular alterations in human tissue. Our goal is to integrate multimodal data and co-register the resected tissues within the whole brain’s *in vivo* MRI space. This approach aims to enhance the characterization and localization of epileptogenic zones and refine surgical treatment targets by identifying abnormalities in global connectivity and structural patterns.

**Significance:** We have successfully developed a systematic protocol for the collection and analysis of multimodal data. This protocol aims to elucidate the structural, functional, and molecular characteristics that render tissue epileptogenic, thereby enhancing the diagnosis and subsequent care of patients with epilepsy.

**Key points:**

1. Introducing a systematic sample handling protocol to assess tissue epileptogenicity at structural, functional and molecular levels.
2. A multimodal approach integrating advanced technology with detailed characterization of epileptogenic tissue properties to obtain patient-specific data.
3. Correlation of data from preoperative MRI and resected tissue to predict tissue pathology from clinical MRI.

## 1 INTRODUCTION

One of the major challenges in epilepsy research is the heterogeneous nature of human epilepsies. They originate from varying etiologies, including brain injuries such as stroke and traumatic brain injury; other structural origins, for instance, tumors, malformations of cortical development, vascular malformations, and glial scars; or from genetic origins (Blumcke et al., 2017). This heterogeneity in etiology and age of onset implies that different mechanisms can result in epilepsy syndromes, calling for personalized strategies for research and clinical practice (Wirrell et al., 2022).

Epilepsy, in general, is a network disease with many indicative zones, such as the epileptogenic zone, seizure onset zone, symptomatogenic zone, irritative zone, functional deficit zone, and a network view of spatiotemporal zones of ictal dynamics (Jehi, 2018)(Barrios-Martinez et al., 2025). This complexity makes it challenging to determine the specific zone in individual patients for resective surgery, which is crucial for curative treatment and for achieving sustained seizure freedom (Laufs, 2012). Nevertheless, resective surgery has been shown to be the most suitable treatment option to control the seizure burden for patients with drug-resistant focal epilepsy, where its efficacy lies in achieving sustained seizure freedom and improving quality of life (Jehi, 2018).

One of the major shortcomings in diagnosing lesional drug-resistant epilepsy is that structural etiology cannot always be recognized noninvasively through visual inspection of MRI (Muhlhofer et al., 2017). Instead, it often requires histopathological analysis, which can only be performed after resective surgery, biopsy, or the invasive implantation of electrodes for intracranial EEG. The pathology remains negative in MRI in 20-40% of cases with focal drug-resistant epilepsy despite histologically verified structural origin (Yang et al., 2024)(Sanders et al., 2024). Although MRI is effective in detecting tumors, vascular lesions, hippocampal sclerosis, and congenital cortical or other malformations, conventional field strength scanners (≤3 Tesla) and MRI methods lack the sensitivity and specificity needed to identify and differentiate smaller tissue alterations from non-epileptogenic lesions and healthy tissue (van Lanen et al., 2021)(De Ciantis et al., 2016). The implementation of novel MRI sequences, along with the detailed characterization of structural, functional, and molecular changes occurring in the tissue, is essential for understanding why tissue becomes epileptogenic.

Earlier studies have made significant efforts to integrate clinical and research teams to explore aspects of tissue epileptogenicity (Straehle et al., 2023) (Andrews et al., 2024)(Buchin et al., 2022). Whereas these studies contribute to understanding DNA methylation, gene expression profile (Kobow et al., 2019), cellular electrophysiology (Wierda et al., 2024), ultrastructure (Alonso-Nanclares et al., 2011), or lesion detection (Spitzer et al., 2022); however, these changes do not occur in isolation; multiple aspects of epileptogenic development must be studied within the same individuals using a protocol that integrates structural, functional, and molecular properties of epileptogenic tissue.

Here, we present the standard operating procedure (SOP) for the multimodal and multiscale assessment of epileptogenic tissue. This procedure was developed through a collaboration involving the clinical epilepsy center, neuroimaging, neurosurgery, and neuropathology departments at Kuopio University Hospital (KUH) in Finland, alongside five multidisciplinary research groups from the A. I. Virtanen Institute for Molecular Sciences (AIVI) at the University of Eastern Finland (UEF).

## 2 METHODOLOGY

### 2.1. Patient recruitment

Patients with drug-resistant focal epilepsy are recruited after detailed multidisciplinary preoperative evaluation and recommendation of epilepsy surgery at the Epilepsy Center at KUH (Kälviäinen et al., 2025). The Ethics Committee of KUH has approved the study for patients in 2015 (128/2015, 21.4.2015), and for healthy volunteers in 2023 (134/2023, 23.05.2023). The tenets of the Declaration of Helsinki are followed. We have also updated the consents and patient information in 2018-2019 according to the European Union General Data Protection Regulation (GDPR) and the preoperative research MRI protocol in 2022. The preoperative research MRI protocol has been approved by Finnish Medical Agency (FIMEA) for patients (FIMEA/2022/006104) and healthy volunteers (FIMEA/2023/004206). Each patient or a legal guardian provides written informed consent to study procedures before any procedure is performed. Patient information is pseudonymized and only the patient code is used for identification.

To illustrate the research pipeline and methodologies, we present a case of a patient with a suspicion of temporal lobe epilepsy from preoperative brain MRI who was 30-40 years old at the time of the surgery. T2-weighted imaging revealed hippocampal atrophy, and following anterior temporal lobectomy (ATL), the patient was diagnosed with hippocampal sclerosis (ILAE type 1) in the right temporal lobe.

### 2.2. Research pipeline

Figure 1 illustrates the data collection workflow. Before resective surgery, patients undergo MRI, including various advanced pulse sequences sequences and a reference sequence for neuronavigation. During surgery, the epileptogenic zone is resected, and its location and orientation are marked with metal clips, which are later tagged in the neuronavigation system. After resection, the entire tissue is immediately placed under oxygenation in ice-cold artificial cerebrospinal fluid (aCSF). The sample is measured and photographed while submerged in oxygenated aCSF. A neuropathologist then slices the tissue in the coronal plane to allocate portions for standard diagnostics and research. Two adjacent slices are reserved for studying tissue microstructure, cellular heterogeneity, and electrophysiological properties at AIVI.

**Figure 1.**
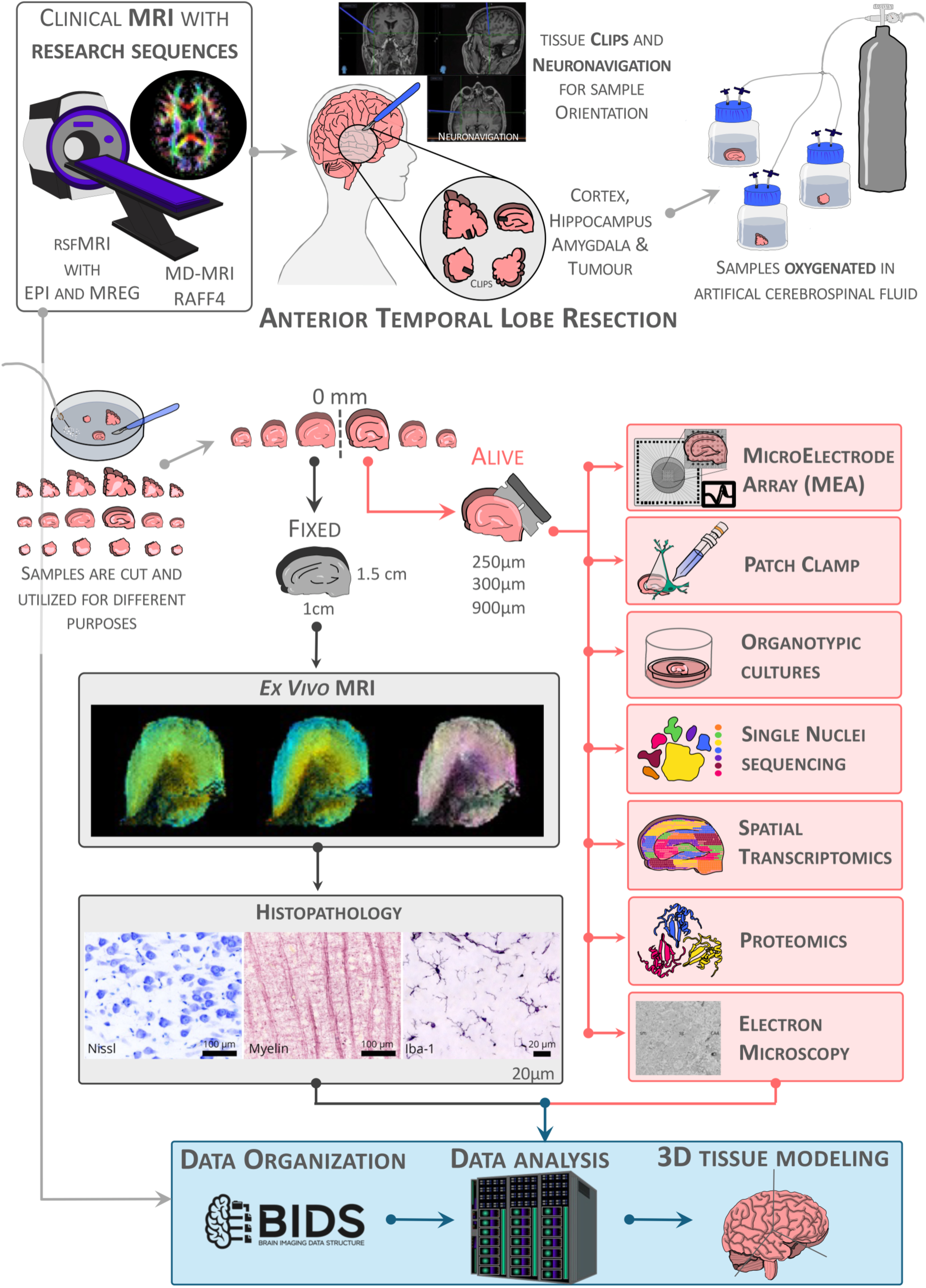
Research pipeline for tissue and data collection. Patients diagnosed with drug-resistant epilepsy are recruited from the Epilepsy Center at Kuopio University Hospital. Five clinical and research groups with complementary expertise collaborate to study the same individuals, from preoperative MRI to the analysis of resected brain tissue. The tissue is transported to the A.I. Virtanen Institute for Molecular Sciences at the University of Eastern Finland, where we investigate its properties using microelectrode arrays (MEA), patch-clamp recordings, organotypic cultures, single-nuclei sequencing, spatial transcriptomics, proteomics, electron microscopy, *ex vivo* MRI, and histology. Ultimately, we integrate preoperative MRI findings with multimodal analyses of the resected tissue to develop highly sensitive MRI approaches for predicting tissue pathology from clinical preoperative imaging.

The midpoint between two consecutive slices, referred to as the “0 mm” surface, serves as the reference for the location and orientation of subsequent procedures. One tissue slice is embedded in low-melting-point agarose gel and sectioned using a vibratome in oxygenated aCSF. Re-slicing from the 0-mm reference surface proceeds as follows: 250-µm thick slices for organotypic cultures, 990-µm slices for immunohistochemistry (IHC), and 300-µm slices for microelectrode array recordings, patch-clamp experiments, single-nuclei sequencing, spatial transcriptomics, proteomics, and 3D-electron microscopy (3D-EM). The second tissue slice is immediately fixed in 4% paraformaldehyde (PFA) and imaged *ex vivo* using an ultra-high-field MRI scanner. Following imaging, this sample is frozen using liquid nitrogen and re-sliced coronally into 20-µm thick sections using a cryostat starting from the 0-mm reference surface for histological analysis.

## 3 RESULTS

### 3.1 Data collection

As of 30.03.2025, we have recruited 26 patients with temporal or extratemporal epilepsy who underwent various surgical procedures, including anterior temporal lobectomy (ATL, n=11), lesionectomy (n=8), combined ATL and lesionectomy (n=2), encephalocele disconnection (n=3), and cortectomy (n=2). Preoperative MRI was acquired from 35 healthy volunteers and 16 of the recruited patients who provided informed consent and met the age criteria for research MRI (18 years or older). Tissue samples were obtained during resective surgery from the cortex, hippocampus, amygdala, or tumor in 20 of the 26 recruited patients.

### 3.2 Preoperative MRI

The current clinical 3T MRI epilepsy protocol includes a set of imaging sequences to assess brain structure and pathology, including T1-weighted 3D-MPRAGE, T2-weighted 3D-FLAIR, susceptibility-weighted imaging, diffusion tensor imaging, and double inversion recovery sequences, with contrast agent administered when necessary (Bernasconi et al., 2019). Our research protocol incorporates novel MRI sequences to enhance sensitivity and specificity in detecting tissue damage, pathology, and functional connectivity. These beyond-state-of-the-art techniques include rotating frame relaxation methods, multidimensional diffusion-relaxation MRI (MD-MRI), and ultrafast functional MRI (fMRI) (Fig. 2).

**Figure 2.**
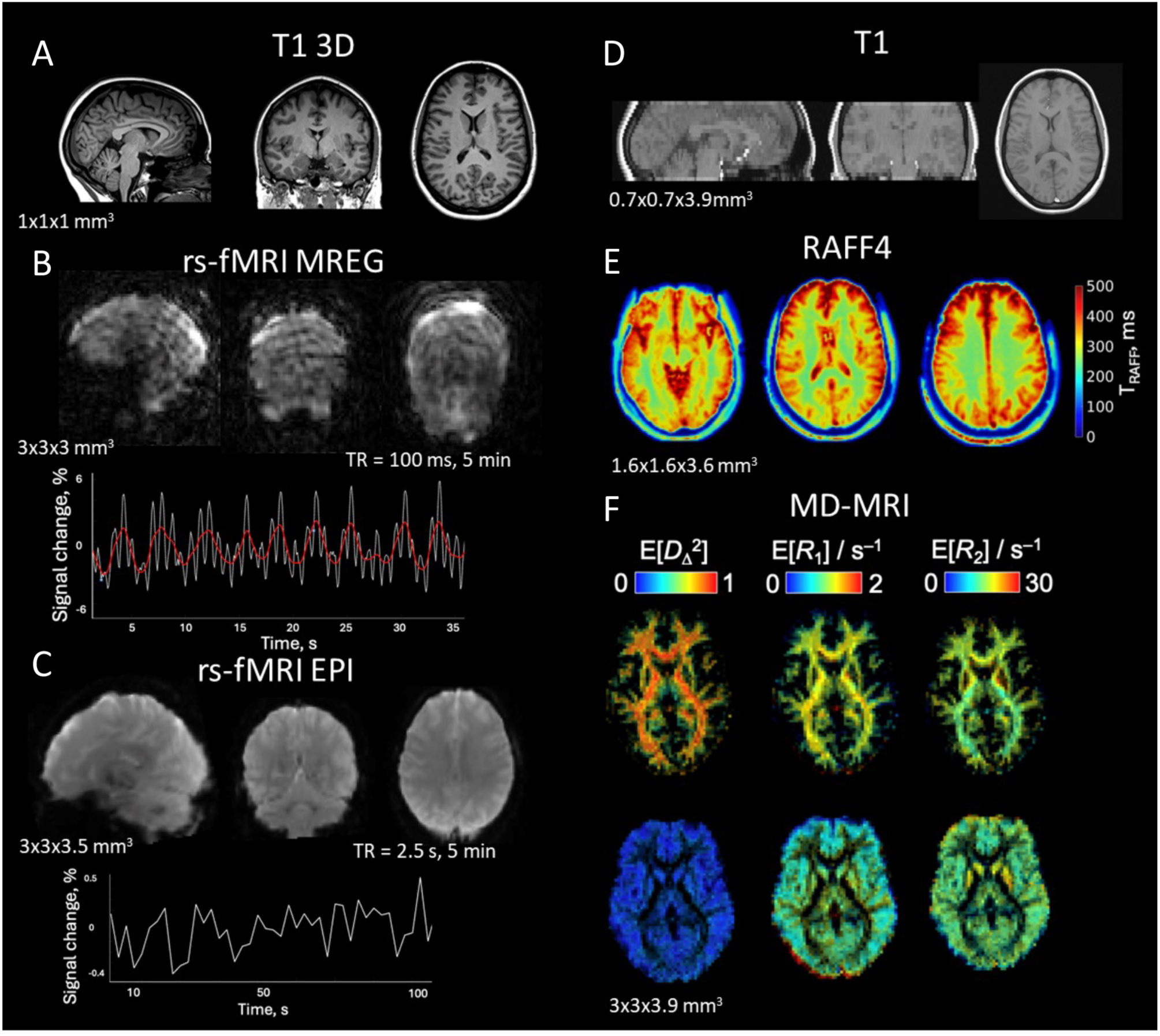
Magnetic resonance imaging (MRI) protocol with research sequences for presurgical imaging. **(A)** T1 weighted 3D image covering the whole brain. **(B)** Magnetic resonance encephalography (MREG) images are shown along with time series from a region of interest inside the brain. White line represents the raw time series, and red line is a smoothed curve. **(C)** Resting-state functional MRI (fMRI) imaged with echo planar imaging (EPI) is shown together with time series from a region of interest. **(D)** T1-weighted reference image shows brain coverage for **(E)** relaxation along the fictitious field 4 (RAFF4) and **(F)** multidimensional MRI (MD-MRI) sequences. **(E)** Relaxation time constant map of RAFF4. **(F)** Relaxation rates parameter maps derived from MD-MRI for white matter and gray matter fractions.

Preoperative imaging is conducted on a 3T Siemens Magnetom Vida MRI scanner (60/200 XT gradients) at KUH approximately 18 hours before the resective surgery. The entire scanning session lasts for 75 minutes. The imaging session starts using a 64-channel Head/Neck Siemens coil. First, we acquire 3D-MPRAGE T1-weighted images (TR = 2.3 s, TE = 1.92 ms, voxel size = 1.0×1.0×1.0 mm^3^), followed by ultrafast fMRI. For that, we utilize magnetic resonance encephalography (MREG), which offers a temporal resolution of 100 ms to direct measurement of breathing- and heartbeat-related fluctuations without aliasing into lower frequencies (Assländer et al., 2013). MREG has proven effective in detecting changes in spectral entropy in drug-resistant epilepsy (Kananen et al., 2020). Our protocol includes a 5-minute resting-state MREG scan (TR = 0.1 s, voxel size = 3.0×3.0×3.0 mm^3^). During the measurement, the patient is instructed to look at the cross displayed on the screen to avoid falling asleep. Heart rate and breathing are recorded during the measurement.

After MREG, the coil is changed to a 20-channel Head/Neck Siemens coil, and the imaging continues by acquiring MD-MRI (slice thickness = 3.0 mm, slice gap = 0.9 mm, 20 slices, in-plane resolution = 3.0×3.0 mm²). MD-MRI is a next-generation multidimensional diffusion-relaxation correlation technique that provides enhanced characterization of heterogeneous microstructures at the voxel level. This method uniquely distinguishes intravoxel size and shape heterogeneity among different cellular populations (Narvaez et al., 2022)(Narvaez et al., 2024). After MD-MRI, we acquired 2D-T1-weighted anatomical images (slice thickness = 3.0 mm, slice gap = 0.9 mm, 20 slices, in-plane resolution = 3.0×3.0 mm^2^) for coregistration purposes. Then, the imaging session continues with relaxation-based MRI sequence relaxation along the fictitious field 4 (RAFF4) (slice thickness = 3.6 mm, 24 slices, in-plane resolution = 1.6×1.6 mm^2^). RAFF leverages a fictitious field in a higher-order frame of reference compared to conventional techniques, enabling specificity across a broader range of spin dynamics while maintaining a clinically acceptable specific absorption rate (Hakkarainen et al., 2016). Notably, RAFF4 has demonstrated superior differentiation between myelination changes and inflammation relative to conventional MRI (Lehto et al., 2017).

In the end of the research imaging session, we perform a 5-minute resting-state fMRI scan with traditional echo-planar imaging (EPI) sequence to map functional brain networks (TR = 2.5 s, TE = 30 ms, slice thickness = 3.5 mm, 42 slices, in-plane resolution = 3.0×3.0 mm^2^). Again, the patient is instructed to keep eyes open and look at the cross.

After that, for patients scheduled for resective surgery, we acquire 3D-T1 MPRAGE (TR = 2.3 s, TE = 1.92 ms, voxel size = 1.0×1.0×1.0 mm^3^) with contrast agent for neuronavigation.

### 3.3 Epilepsy surgery at KUH and tissue delivery to AIVI/UEF

Representatives from the research team join the clinical team in the operating room (OR) at the time of the surgery. They document key details regarding the location and orientation of the resected tissue. In the case of a temporal lobe resection, metal clips are affixed to anatomically defined regions of the resected tissue to facilitate precise anatomical landmarks required for subsequent analyses: 1) posteriorly to the anterior temporal gyri, 2) anteriorly to the hippocampus, and 3) laterally to the amygdala. The T1-weighted anatomical neuronavigation images are integrated into the neuronavigation system and used by the surgical team for registering the location of metal clips. The coordinates of the clips are saved by the system and exported to a JSON file to enable specimen localization and volumetric re-construction.

The resected tissue is immediately placed into pre-oxygenated ice cold aCSF to preserve the tissue viability and connected to continuous oxygenation immediately. Then, the tissue is sectioned by a neuropathologist, still under oxygenation. The tissue slices are divided for diagnostic histopathology and the two research teams at AIVI.

### 3.4 Brain slice preparation for electrophysiological experiments

One of the tissue slices resected during the surgery goes immediately under oxygenation to the neurophysiology laboratory. The tissue is gently cleaned of any debris or blood clots and photographed for sample size documentation. The slice is then embedded in 2% TopVision low melting point agarose (Thermo Fisher Scientific) and sliced (250 µm for organotypic cultures, 990 µm IHC and spatial transcriptomics, and 300 µm for microelectrode array (MEA) recordings, patch-clamp experiments, single-nuclei sequencing, proteomics, and 3D-EM) using a vibratome (Campden Instruments) in chilled (2-4°C), fully carbogenated (95%/5%, O_2_/CO_2_) aCSF of the following composition (mM): 92 NMDG-methyl-D-glucamine, 2.5 KCl, 20 HEPES, 25 NaHCO_3_, 1.25 NaH_2_PO_4_, 3 Na-pyruvate, 2 thiourea, 5 Na-ascorbate, 7 MgCl_2_, 0.5 CaCl_2_, 25 glucose (pH = 7.3). After cutting, slices are placed in a custom-made chamber and allowed first to recover at 34°C for 45 minutes in the recording solution (see 3.6) supplemented with 3 mM Na-pyruvate, 2 mM Thiourea and 5 mM Na-ascorbate) and then for another 60 minutes in the same solution at room temperature (20-22°C) before electrophysiology.

### 3.5 3D Microelectrode array (MEA)

MEA recordings can monitor local field potential (LFP) occurrence and synchronization (Rigas et al., 2015), multiunit spiking behavior (Tóth et al., 2018), emergence of stimulus/drug-induced multiband oscillations (Carmeli et al., 2013) and network connectivity (Dastgheyb et al., 2020). A single slice of 300 µm is positioned into a 60-electrode 3D MEA (Multichannel Systems, MCS) and secured with an anchor. Signals, such as spike and LFP with or without 4-aminopyridine (4-AP), a potassium channel blocker used to mimic epileptogenic activity, are obtained with an MEA2100-Mini-system (MCS) (Fig. 3). Recordings are performed under constant perfusion (3 ml/min) with fully oxygenated (95% O_2_/5% CO_2_) aCSF at 32-34°C (Fagerlund et al., 2022). Custom and planar electrode MEAs are available as well if required for specific research questions during the experiments.

**Figure 3.**
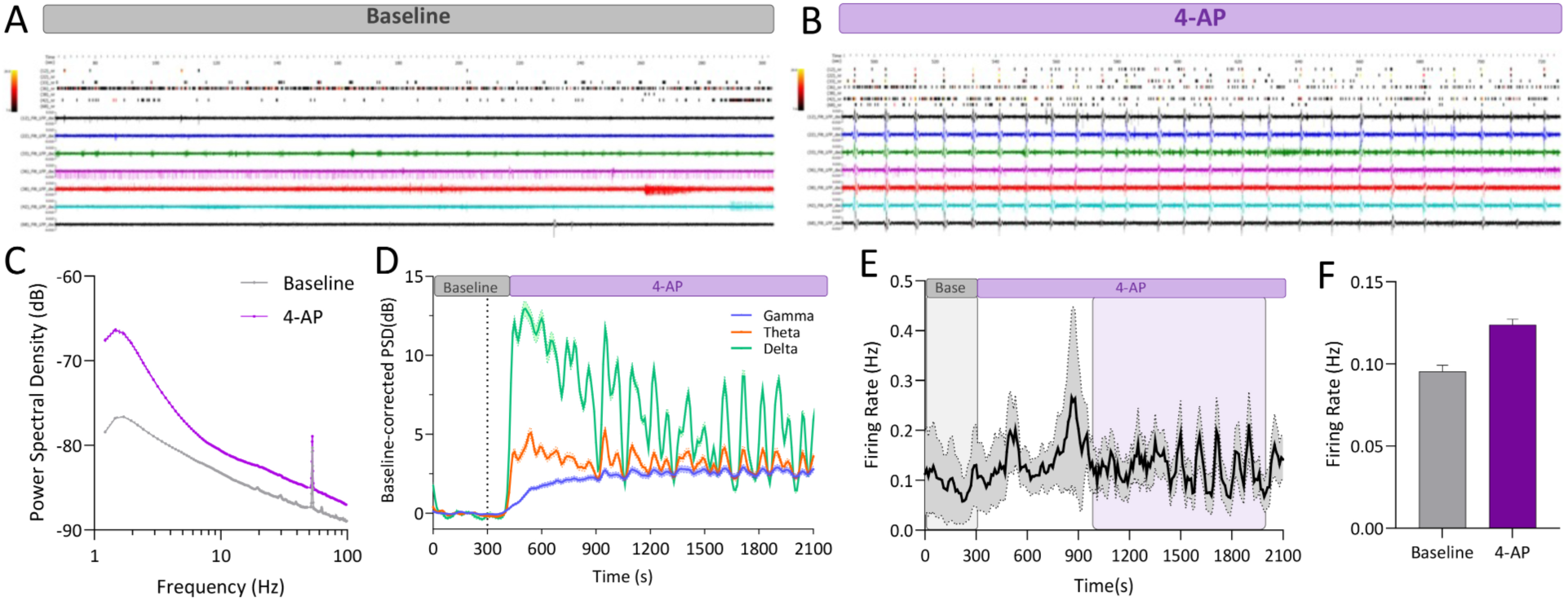
Spike and LFP results from cortical epilepsy sample treated with 4-aminopyridine (4-AP), a potassium channel blocker used to mimic epileptogenic activity. Raster plots depicting spike and LFP data from top seven spiking electrodes (more than 1000 timestamps) from baseline **(A)** and treated with 4-AP **(B)**. Upon 4-AP there is a clear enhancement of network activity and LFP-spike time locked events. **(C)** Baseline-corrected gamma, theta and delta power spectral densities (PSD) against time depicting the baseline period and exposure to 4-AP. **(D)** Power Spectral Density (dB) against time for the baseline period and exposure to 4-AP. There is a clear difference in all the frequency spectrum when 4-AP is applied. **(E)** Firing rate timeline depicting the baseline period and exposure to 4-AP. The coloured boxes indicate the time period from which the averaged firing rate was calculated and depicted as a boxplot **(F)**. Only channels with a mean firing rate higher than 0.05 spikes/s are considered in the analysis.

### 3.6 Whole-cell patch-clamp recordings and neuromorhometric reconstruction

Single slices from the cortex and hippocampus (300 μm) are transferred and secured with a slice anchor in a large volume bath under an Olympus BX50WI microscope equipped with differential interference contrast (DIC) optics, a 40x water immersion objective and a CCD camera (Retiga R1, Q-imaging). Slices are continuously perfused at a rate of 2.5-3 ml/min with a recording solution of the following composition (mM): 120 NaCl, 2.5 KCl, 25 NaHCO_3_, 1.25 NaH_2_PO_4_, 2 CaCl_2_, 1 MgCl_2_, 25 glucose. To induce epileptiform activity 4-AP (100 μM, (Luhmann et al., 2000)) is added to recording solution for long-term recording of spontaneous activity. Whole-cell current and voltage-clamp recordings were conducted at 32-33°C with an Axopatch-200B amplifier (Molecular Devices) using 5-8 MΩ glass electrodes filled with an internal solution containing (mM): 135 potassium-gluconate, 5 NaCl, 10 HEPES, 2 MgCl_2_, 1 EGTA, 2 Mg-ATP, 0.25 Na-GTP and 0.5% Biocytin (pH adjusted to 7.3 with osmolarity at 275-285 mOsm/l). Electrophysiological data are low pass filtered at 1 KHz (4-pole Bessel filter) then captured at 10 KHz via a Digidata 1440A A/D board to a personal computer, displayed in Clampex software and stored to disk for further analysis. Excitatory and inhibitory spontaneous synaptic current (EPSCs and IPSCs) are sampled at a holding potential of −70 mV and to −40 mV (to avoid recruitment of action potentials) respectively in 2-minute epochs as reported previously (Dougalis et al., 2012). To examine current-input spike-output firing relationships and subthreshold currents, the neurons are given, 1s incremental current injections of ± 5-50 pA from their resting membrane potential to see the AP firing pattern correspond to specific neuronal cell type (Lee et al., 2023).

Whole patch-clamp recordings are performed in a cortical layer 2/3 (up to 0.25-1.2 mm depth from pia). Under the long-term application of 4-AP (100 μM) (Luhmann et al., 2000) the excitatory and inhibitory neurons of the cortex and hippocampus exhibit the features of epileptic tissue, such as the appearance of giant depolarizing currents, synchronization and increased number of action potentials. Biocytin filling of patched neurons is done to track the morphological features (e.g., total dendritic length, total nodes, spine density) of the recorded cells as described in (Dougalis et al., 2025). Pyramidal neurons and intraneuronal cells are discriminated by their exhibited active membrane properties (firing pattern) (Fig. 4A**-C**) and by the following post-hoc *in situ* morphological characteristics of the dendritic tree.

**Figure 4.**
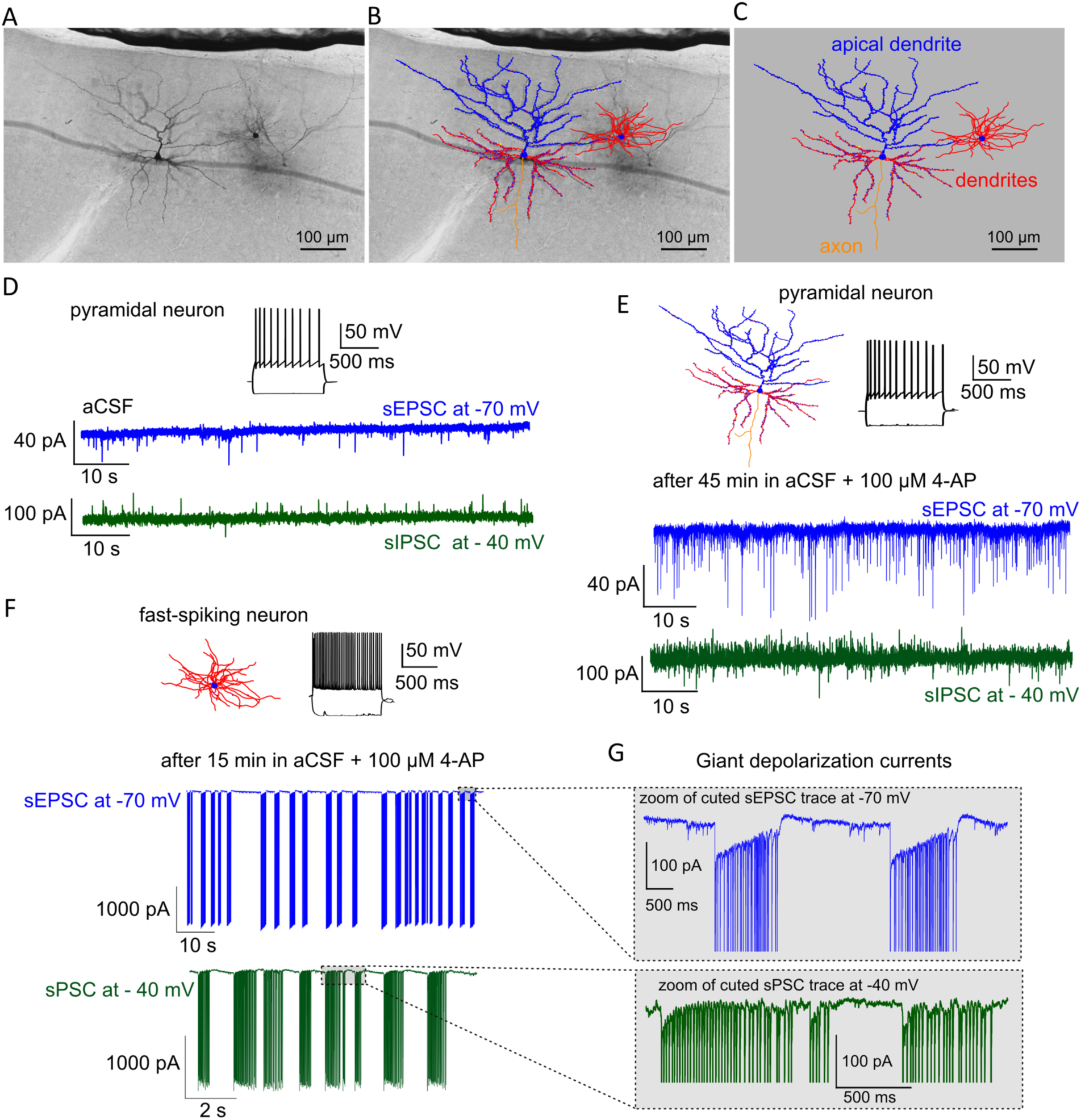
Patch clamp recording from a right temporal cortex slice. **(A)** Brightfield image of patched neurons filled with biocytin. **(B)** Morphology reconstruction of pyramidal neuron and fast-spiking interneuron expressed in cortical layer 2/3 overlay on brightfield microscopy image. **(C)** Neuromorphometric reconstruction of biocytin-filled neurons with distribution of basal dendrites (red), apical dendrite (blue) and axon (yellow). **(D)** AP firing pattern of pyramidal neuron recorded in aCSF and their spontaneous excitatory postsynaptic currents (sEPSC) recorded at voltage clamped at −70 mV, and spontaneous inhibitory postsynaptic currents (sIPSC) recorded at −40 mV. **(E)** Neuronal morphology reconstruction and AP firing pattern of the pyramidal neuron recorded after 45 min action of 100 μM 4-AP in aCSF, and their sEPSC recorded at −70 mV and sIPSC recorded at −40 mV. **(F)** Neuronal morphology reconstruction and AP firing pattern of the fast-spiking interneuron recorded after 15 min action of 100 μM 4-AP in aCSF, and their epileptiform activity presenting as giant sEPSC recorded at −70 mV and this epileptiform activity recorded at −40 mV. **(G)** Zoom of events are presented as giant postsynaptic currents.

Biocytin-filled patch-clamped neurons from recorded slices were subjected to DAB staining and neuromorphometric reconstructions (Mohan et al., 2015). Briefly, slices are fixed in fresh 4% PFA for 48-72 hours at 4°C and then are stored indefinitely in a phosphate buffered saline (PBS) with 0.1% sodium azide at 4°C until processed. Reconstructions are performed with a calibrated XYZ axis Ludl electronics stage under an x100 oil objective of a Nikon Eclipse microscope using Neurolucida software (MBF Biosciences). For reconstructions, we select only neurons where the apical and the basal trees were largely preserved intact. Analysis of the morphological features (e.g. total dendritic length, total nodes, spine density) for neurons are conducted via Neurolucida Explorer software.

### 3.7 Organotypic cultures

Organotypic slice cultures (OCs), *i.e.* cultures of tissue over extended period of time, enable measurement of tissue electrophysiological properties over time and offer a platform for testing different therapeutic approaches. The OCs are cultured as described by (Bak et al., 2024) with small modifications. The tissue is sectioned into 250-µm slices and cultured on 6-well plate inserts. During the first hours, they are stabilized in cell culture media with a cocktail of cytokines and later transferred to human cerebrospinal fluid. Electrophysiological activity is recorded using an MEA system (see 3.5) and induced with perfusion of 1 mM 4-AP to block voltage-sensitive K+ channels or 200 µM N-methyl-D-aspartate (NMDA) to activate NMDA receptors. Adeno-associated virus vectors are used to quantify the viable neurons and specific neuronal types.

### 3.8 Single-nuclei sequencing

Single nucleus RNA sequencing (snRNA-seq) enables a deeper understanding of each cell’s diversity of function and dysfunction in the context of epilepsy. Recent snRNA-seq studies from resected tissues have pinpointed specific dysregulations in glutamate signaling (Pfisterer et al., 2020), reactive astrocytes (Pai et al., 2022) and microglial activation (Kumar et al., 2022). However, how altered cell states contribute to neuronal hyperexcitability in electrogenesis is understudied. To isolate nuclei from fresh-frozen samples, we use a previously published protocol (Gazestani et al., 2023) followed by fluorescence-activated nuclei sorting for DAPI+nuclei selection and snRNA-seq. Complementary DNA libraries are then prepared using the Chromium GEM-X Single Cell 3’ Kit v4 (10x Genomics) following the manufacturer’s protocol. Libraries are sequenced on Novaseq X platform (Illumina). Cell types of each cluster are annotated by cell type specific signatures. The candidate disease drivers are identified, and the individual gene transcription profiles are quantitatively compared between the samples. Cellular pathways are analyzed in relation to epilepsy, highlighting different stages of microglial activation and distinct neuronal subpopulations. (Stuart et al., 2019).

### 3.9 Spatial transcriptomics

To investigate how the altered cellular states are spatially organized, we conduct spatial transcriptomics analysis with Xenium Analyzer using Xenium Prime 5K pan tissue and pathways panel (all 10x Genomics) following the manufacturer’s protocol. The analysis allows transcript identification with spatial mapping of various cellular subpopulations in relation to the epilepsy-related pathology across tissue section. We map altered cell populations from the snRNA-seq dataset onto the spatial resolution of spatial transcriptomics creating a dataset with spatially resolved high-resolution cell-specific transcriptomic profiles. First, we annotate cell populations in both datasets, then apply a transfer learning algorithm to assign spatial coordinates to cell populations in the snRNA-seq data. This approach overcomes the limitations of each technology (no spatial resolution in snRNA-seq and few transcripts per cell in spatial transcriptomics) and was previously implemented in study by (Scoyni et al., 2024).

### 3.10 Proteomics

Untargeted proteomics is used for proteome profiling. The tissue section allocated for proteomics (300 µm) is homogenized and enzymatically digested, peptides are separated by liquid chromatography and quantified by tandem mass spectrometry. Additionally, activity-based protein profiling is used to measure enzymes activity in the same sample. Part of endocannabinoid system, which is linked to increased neuronal excitability related to human epilepsy, is studied using markers for fatty-acid amide hydrolase 1 (FAAH) and monoacylglycerol lipase (MAGL) (Romigi et al., 2010).

### 3.11 3D-electron microscopy (3D-EM)

Recent advancements in EM allow 3D imaging of the tissue and larger field of view of tissue (Peddie et al., 2022). Here, 300-µm thick sections (see 3.4) are immediately immersed in a solution of 2% PFA/2.5% glutaraldehyde overnight at 4°C and then placed in 2% PFA in 0.15 M cacodylate buffer and 2 mM CaCl_2_ (pH = 7.4) until staining (Fig. 5J). The sections are stained using an enhanced staining protocol (Fig. 5K) and imaged using serial block face scanning electron microscopy (Salo et al., 2021). After selecting the area of interest for imaging, the blocks are further trimmed and mounted using silver epoxy (CircuitWorks CW2400) on aluminum specimen pins, painted with silver paint and sputtered with a 5-nm layer of platinum coating (Q150TS coater, Quorum Technologies, Laughton, UK).

**Figure 5.**
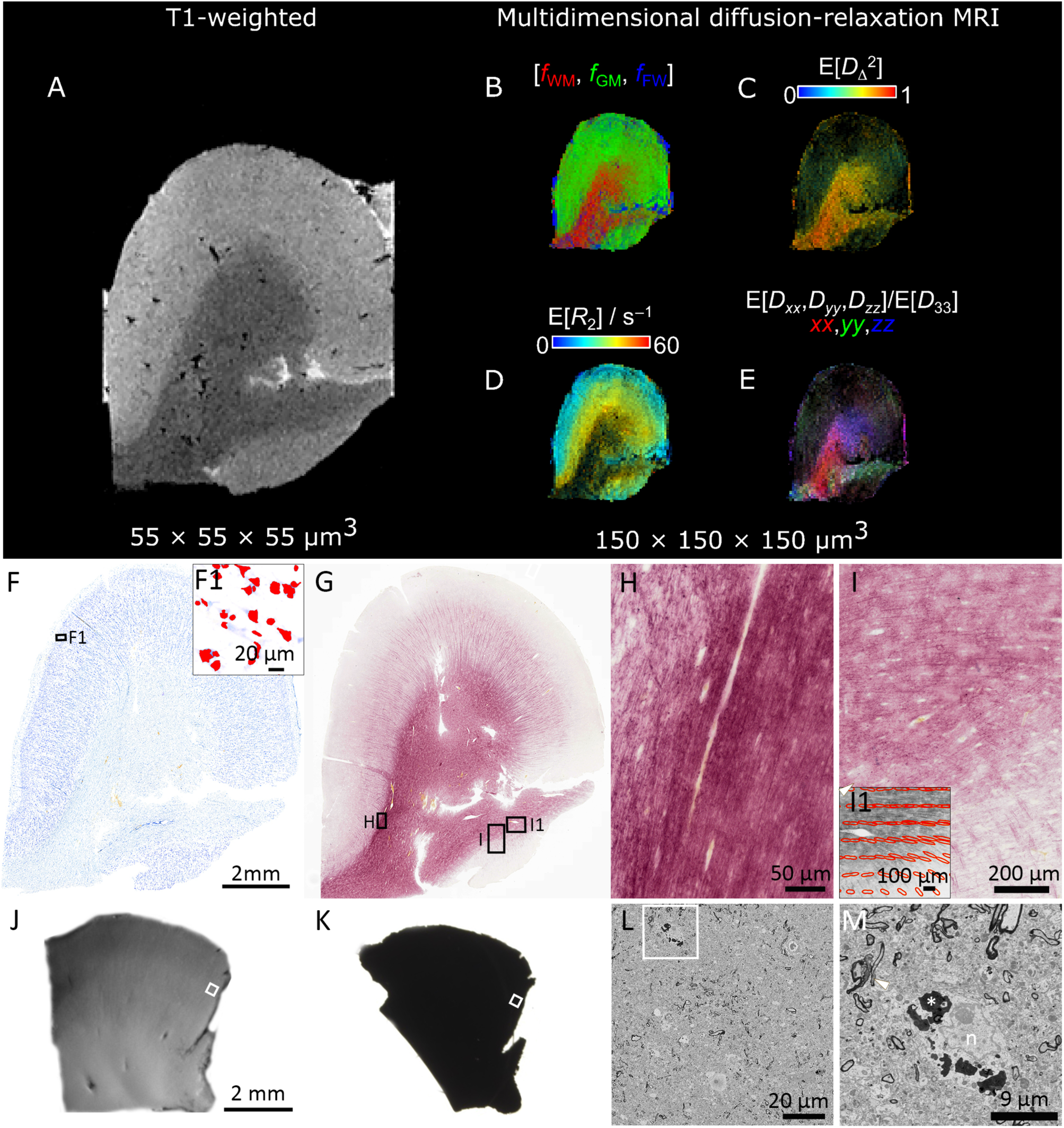
*Ex vivo* MRI and histology from the right temporal cortex. **(A)** T1 weighted images with a voxel size of 55 x 55 x 55 µm^3^. **(B-E)** Multidimensional diffusion-relaxation maps with a voxel size of 150 x 150 x 150 µm^3^. **(B)** A map that shows white matter (WM), grey matter (GM) and phosphate buffered saline (PBS) fractions derived from the diffusion-relaxation distributions. **(C)** The per voxel mean of the squared normalized anisotropy parameter *D*_Δ_^2^ from the WM fraction. **(D)** The transverse relaxation rate *R*_2_ for the GM fraction. **(E)** The average orientation of the diffusivity for the WM fraction, highlighting the orientation of the axons bundles. **(F)** A representative photomicrograph showing overall cytoarchitecture of Nissl-stained section. **(F1)** Zoom in into the cortical layer II and cells segmented with automated cell detection enabling analysis of cell size, morphology and density. **(G)** A representative photomicrograph showing myeloarchitecture of a gold chloride-stained section. Black boxes in panel **G** show higher magnification areas in panels **H** and **I**. **(H) and (I)** Two high-magnification photomicrographs showing the interface between cortical layer VI and WM. Note the similarities as shown in panel **C**. **(I1)** Zoom in into GM and WM interface with structure tensor analysis detecting orientation of the myelinated axons. **(J)** Fixed 300-µm thick section for 3D-EM; **(K)** same section stained using an enhanced staining protocol with heavy metals; **(L)** a representative image of the dataset with a resolution of 15 x 15 x 50 nm^3^. White squares indicate the area of interest for trimming and imaging; and **(M)** a close-up into the same image. Arrowheads indicate myelinated axons, asterisk lipofuscin and n nucleus.

The blocks are imaged on an SEM microscope (Quanta 250 Field Emission Gun; FEI Co., Hillsboro, OR) equipped with 3View system using a backscattered electron detector (Gatan Inc., Pleasanton, CA). For imaging, we use a 2.5-kV beam voltage, spot size 3 and 0.08-0.22 Torr of pressure. We acquire datasets of a volume of 16x16x65 μm^3^ with a voxel size of 15x15x40 nm^3^ (Fig. 5L**-M**); and of a volume of 160x80x65 μm^3^ with a voxel size of 40x40x40 nm^3^. After imaging, we pre-process SBF-SEM image stacks with Microscopy Image Browser (MIB; http://mib.helsinki.fi; (Belevich et al., 2016)), and obtained 2D and 3D microscopy data are quantitatively analyzed by in-house automated and customized methods (Abdollahzadeh et al., 2021), (Salo et al., 2021).

### 3.12 *Ex vivo* imaging

*Ex vivo* MRI provides high-resolution imaging of resected brain tissue, enabling detailed microstructural analysis without motion artifacts. It allows direct correlation with histopathology for validating imaging biomarkers of epileptogenic tissue while preserving spatial context for accurate localization of histological findings.

The resected sample designated for *ex vivo* MRI is fixed in 4% PFA for at least 3 days, then scanned in a vertical 11.7-T MRI scanner. The tissue is immersed in perfluoropolyether (Solexis Galden^Ⓡ^) to avoid signals outside the tissue. The scanning protocol includes T1-weighted (TR = 15 ms, TE = 5 ms, and isometric voxel 55 µm^3^), T2-weighted (TR = 2000 ms, TE = 65 ms, and isometric voxel 55 µm^3^), RAFF4 (TR = 2500 ms, TE = 11.5 ms, and isometric voxel 150 µm^3^), single diffusion encoding (SDE) (TR = 900 ms, TE = 34 ms, 3 b-values and isometric voxel 100 µm^3^) and MD-MRI (variable TR, TE, b-values, b-tensor encoding, and centroid frequency with isometric voxel 150 µm^3^). RAFF4 and SDE are acquired using spin echo-echo planar imaging (SE-EPI) and MD-MRI data with SE-EPI modified with free gradient waveforms.

MD-MRI utilizes frequency-dependent tensor-valued diffusion MRI correlated with longitudinal and transverse relaxation rates expressed as nonparametric **D**(ω)-*R*_1_-*R*_2_ distributions to provide sub-voxel level information (Narvaez et al., 2022). The distributions are used to separate different water populations within a single voxel by manually binning the parameter space, resulting in white matter, grey matter, and PBS. The objective is to evaluate the diffusion, relaxation and restriction properties given by MD-MRI and validate the signal by performing histological analysis from the same specimen after scanning (Fig. 5). Derived from MD-MRI, per voxel **D**(ω)-*R*_1_-*R*_2_ distributions are obtained that can be divided to highlight tissue fractions (i.e., white matter, grey matter, and PBS) (Fig. 5B). Squared normalized anisotropy (*D*_Δ_^2^) from the white matter fraction provides information about the axon integrity (Fig. 5C). Transverse relaxation rate (*R*_2_) of the grey matter fraction displays the myelinated axons in cortical layers (Fig. 5D). The diffusion orientation from the white matter fraction also reveals principal axon bundle directions (Fig. 5E).

### 3.13 Histology

Histology provides detailed cellular and tissue-level information essential for understanding the structural alterations associated with epilepsy. By incorporating quantitative analyses, we can extract tissue metrics such as neuronal density, inflammation, myelin alterations, synaptic organization, and other features that contribute to epileptogenesis. These analyses help differentiate pathological subtypes, validate imaging biomarkers, and establish correlations between molecular, electrophysiological, and structural changes. In our pipeline, we perform histology after *ex vivo* MRI (3.12), after MEA measurements (3.5 and 3.7) and in sections designated for IHC (3.4). MEA and IHC sections are fixed with 4% PFA after measurements and sectioning, respectively. Tissue from all these three modalities is cryoprotected with 30% sucrose in phosphate buffer, frozen on liquid nitrogen and re-sliced into 20-µm sectioned using cryostat.

For MRI validation, we use systematic histopathological and quantitative approach to understand the tissue microstructure underlaying MRI. We use Nissl staining (thionin) to study the overall cytoarchitecture (Fig. 5F**-I**) and myeloarchitecture (Fig. 5F**-I**). Immunostaining is performed to detect changes in neuronal morphology (neuronal nuclear antigen, NeuN) and glial cells as markers of inflammation (ionized calcium-binding adaptor molecule 1 (Iba1) and glial fibrillary acidic protein (GFAP)). After digitalization with a slide scanner (Olympus), stained sections are quantified for cell and myelin density, sole analysis or structure tensor from individual cortical layers and hippocampal subfields (Fig. 5**F1, I1**). Sections MEA measurements and exclusively designated for IHC are re-sliced and are stained with Nissl, neuronal (NeuN), microglia (Iba1) and astrocyte (GFAP) markers as well as makers for cytoskeleton (microtubule-associated protein 2 (MAP2)), synapses (synapsin1) and others. High-resolution microscopy with 3D reconstruction of glia-neuron interactions is carried out for evaluation of synaptic loss and spine engulfment using synapsin 1 and NeuN antibodies.

### 3.14 Multimodal co-registration of resected tissues in the whole brain MRI

Predicting the signature patterns of epileptogenic tissue requires implementation of dedicated algorithms to model the relationships between multiscale imaging, electrophysiological, cellular, and histological data. As the first step for multiscale data integration, Fig. 6 demonstrates the layout of Medtronic neuronavigation system in which the precise location of inserted metal clip in the anterior temporal lobe is intraoperatively registered by the surgeon. Next, the coordinates from this marker are used to localize the resected cortical specimen on preoperative research-sequence 3D-T1-weighted and its 3D reconstructed pial surface. Finally, the exact location and orientation of resected specimen are obtained with the help of intraoperative notes and morphology of gyri and sulci in the resected tissue. This step helps in identification of epileptogenic zones in preoperative MRI sequences for precise surgical and treatment planning.

**Figure 6.**
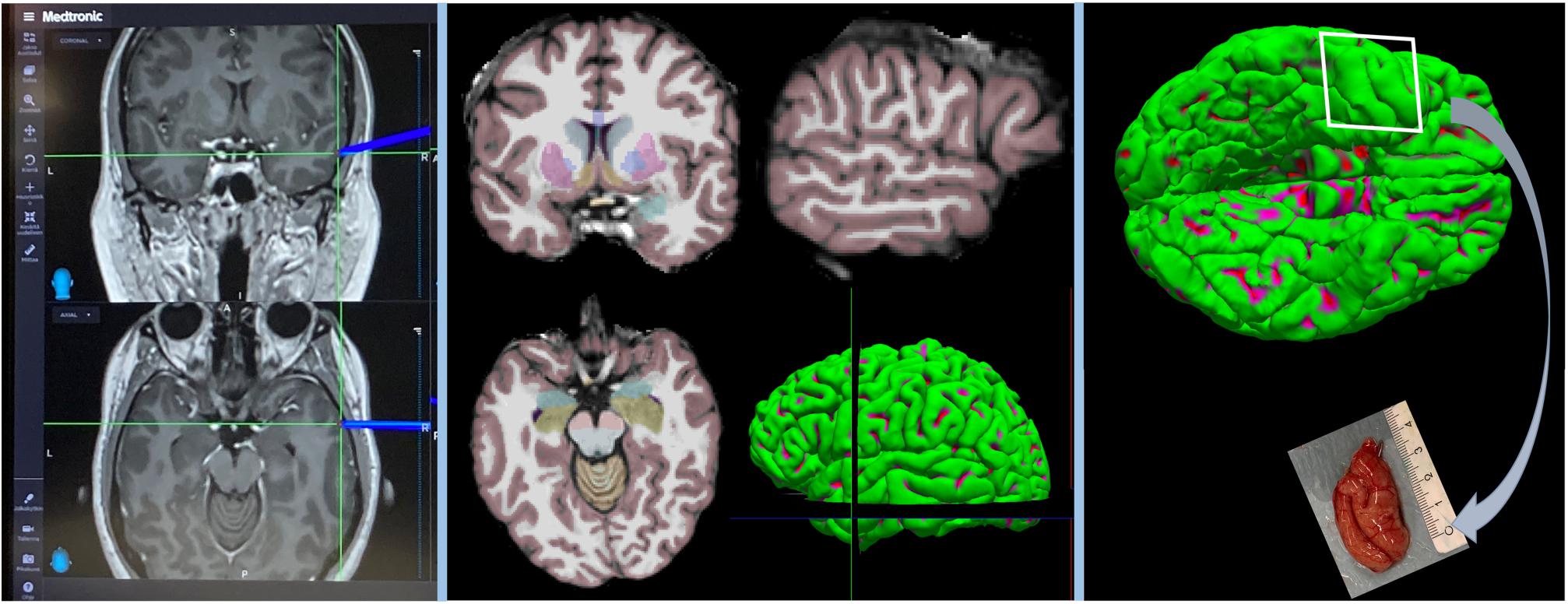
Multiscale data integration as a bridge between pre-operative *in vivo* MRI to postoperative localization of resected specimen. **(A)** Neuronavigation-guided registration of metal clip inserted to the posterior surface of a cortical specimen on the MRI space. **(B)** Localization of registered marker in the coronal, sagittal, and axial slices of the 3D T1w research-sequence, pre-operative MRI. Also, FreeSurfer-based cortical and subcortical segmentation and volumetric reconstruction of pial surfaces are shown. **(C)** Location and orientation of resected cortical specimen according to morphometry (gyri and sulci shape) in tip of the anterior temporal lobe.

### 3.15 Documentation and data organization

The intact, segmented and sliced samples are photographed from all surfaces to enable reconstruction. The researcher(s) who enter the OR record the events and their timestamps from preoperative phase to post-operative tissue dissection and compile them into a machine-readable spreadsheet. One of the aims is to introduce a research data management (RDM) system for integration of sensitive patient data and research protocols. This system adheres to the Findable, Accessible, Interoperable, and Reusable (FAIR) principles (Wilkinson et al., 2016) and enables efficient and responsible utilization of multimodal and multiscale data from dissected tissues in a project on drug-resistant epilepsy. The implemented RDM system encompasses data storage, computation, documentation, and workflow management. Research data is securely maintained on an access-controlled disk system connected to high-performance computing resources. Data organization is compliant with the Brain Imaging Data Structure (BIDS) neuroimaging framework (Gorgolewski et al., 2016). Customized data organization and protection plans ensure the safe handling of patient data in line with the European and Finnish regulations. Documentation includes version-controlled Sample Operating Procedures (SOP) tailored to epilepsy surgeries such as anterior temporal lobectomy, lesionectomy, encephalocele, and disconnection. Finally, workflow management facilitates adherence to SOPs by streamlining communication between clinical and research teams.

## 4 DISCUSSION

As a combined effort between five clinical and research groups at the KUH and UEF, we have developed systematic sample handling and multimodal analysis protocol to assess tissue epileptogenicity at structural, functional and molecular levels.

By introducing advanced preoperative MRI methods, we expect improvement in the sensitivity and/or specificity to identify and differentiate smaller tissue alterations associated with epileptogenicity. Furthermore, we can explore functional local network changes and alteration in brain pulsations that modulate large-scale networks generating spontaneous seizures and epilepsy. Finally, we can use the information to enhance detection of location of epileptogenic zone and refine targets for surgical treatment based on abnormalities in global connectivity and structural patterns. *Ex vivo* MRI with ultra-high resolution provides a bridge between preoperative MRI and cellular level changes.

The pathological mechanisms of epilepsy are highly complex, and their understanding is fundamental for early diagnosis, therapy, and prognosis of patients with epilepsy. Having gained understanding of the synaptic signatures and glial activation states at the functional, ultrastructural, and transcriptional state in the brains of patients with epilepsy, we will be able to use this information as readouts for epileptogenic zone detection. Here, we open new avenues for unveiling temporal lobe epilepsy-related pathological mechanisms such as inflammation and neuronal function by multiscale imaging, electrophysiological, cellular, and histological data. This multimodal data can ultimately reveal human neuronal network functions also outside the structural lesion and ultrastructural alterations related to focal structural drug-resistant epilepsy.

Our pipeline is fully operational, yet we continuously refine our methodologies with each patient and sample collection, adapting to variations in tissue availability, partial sample acquisitions, patient consent modifications, and scheduling constraints to maintain flexibility and optimize our research workflow. Future directions focus on enhancing data processing and analysis workflows, optimizing multimodal data integration, and refining quantitative methods for characterizing epileptogenic tissue. Additionally, efforts will be directed toward identifying patient-specific patterns, enabling more precise diagnostics and personalized treatment strategies for epilepsy.

## AUTHOR CONTRIBUTIONS

*Conceptualization:* Olli Gröhn, Reetta Kälviäinen, Alejandra Sierra, Tarja Malm, Jussi Tohka; *Investigation:* Jenni Kyyriäinen, Adriana Della Pietra, Mastaneh Torkamani-Azar, Mireia Gómez Budia, Polina Abushik, Nataliia Novosolova, Henri Eronen, Omar Narvaez, Ekaterina Paasonen, Vera Lezhneva, Anssi Pelkonen, Liudmila Saveleva, Antti Huotarinen, Ilya Belevich, Eija Jokitalo, Kuopio Epilepsy Center Epilepsy Surgery Group, Tuomas Rauramaa, Arto Immonen, Ville Leinonen*; Data Curation:* Jussi Tohka, Mastaneh Torkamani-Azar*; Writing – Original Draft:* Jenni Kyyriäinen, Alejandra Sierra*; Writing – Review & Editing:* Olli Gröhn, Reetta Kälviäinen, Alejandra Sierra, Tarja Malm, Jussi Tohka, Liudmila Saveleva, Mastaneh Torkamani-Azar, Ekaterina Paasonen, Omar Narvaez*; Visualization:* Mireia Gómez Budia, Ekaterina Paasonen, Mastaneh Torkamani-Azar, Polina Abushik, Omar Narvaez, Jenni Kyyriäinen*; Project administration:* Olli Gröhn, Reetta Kälviäinen, Alejandra Sierra, Tarja Malm, Jussi Tohka; *Funding acquisition:* Olli Gröhn, Reetta Kälviäinen, Alejandra Sierra, Tarja Malm, Jussi Tohka.

## ACKNOWLEDGEMENTS

Part of the work was carried out with the support of the *In Vitro* and *Ex Vivo* Electrophysiology core facility (part of Biocenter Kuopio and Biocenter Finland), Bioinformatics Center, University of Eastern Finland, Finland. *Ex vivo* MRI is carried out with the support of Kuopio Biomedical Imaging Unit, University of Eastern Finland, Kuopio, Finland (part of Biocenter Kuopio, Finnish Biomedical Imaging Node, and EuroBioImaging). SBF-SEM is performed at the Electron Microscopy Unit of the Institute of Biotechnology, University of Helsinki supported by Helsinki Institute of Life Science and Biocenter Finland.

## ETHICAL PUBLICATION STATEMENT

We confirm that we have read the Journal’s position on issues involved in ethical publication and affirm that this report is consistent with those guidelines.

## REFERENCES

Abdollahzadeh, A., Belevich, I., Jokitalo, E., Sierra, A., & Tohka, J. (2021). DeepACSON automated segmentation of white matter in 3D electron microscopy. Communications Biology, 4(1), 1–14. 10.1038/s42003-021-01699-w

Alonso-Nanclares, L., Kastanauskaite, A., Rodriguez, J. R., Gonzalez-Soriano, J., & Defelipe, J. (2011). A Stereological Study of Synapse Number in the Epileptic Human Hippocampus. Frontiers in Neuroanatomy, 5(FEB), 8. 10.3389/FNANA.2011.00008

Andrews, J. P., Geng, J., Voitiuk, K., Elliott, M. A. T., Shin, D., Robbins, A., Spaeth, A., Wang, A., Li, L., Solis, D., Keefe, M. G., Sevetson, J. L., Rivera de Jesús, J. A., Donohue, K. C., Larson, H. H., Ehrlich, D., Auguste, K. I., Salama, S., Sohal, V., … Nowakowski, T. J. (2024). Multimodal evaluation of network activity and optogenetic interventions in human hippocampal slices. Nature Neuroscience, 27(12), 2487–2499. 10.1038/s41593-024-01782-5

Assländer, J., Zahneisen, B., Hugger, T., Reisert, M., Lee, H.-L., LeVan, P., & Hennig, J. (2013). Single shot whole brain imaging using spherical stack of spirals trajectories. NeuroImage, 73, 59–70. 10.1016/j.neuroimage.2013.01.065

Bak, A., Koch, H., van Loo, K. M. J., Schmied, K., Gittel, B., Weber, Y., Ort, J., Schwarz, N., Tauber, S. C., Wuttke, T. V, & Delev, D. (2024). Human organotypic brain slice cultures: a detailed and improved protocol for preparation and long-term maintenance. Journal of Neuroscience Methods, 404, 110055. 10.1016/j.jneumeth.2023.110055

Barrios-Martinez, J. V., Almast, A., Lin, I., Youssef, A., Aung, T., Fernandes-Cabral, D., Yeh, F. C., Chang, Y. F., Mettenburg, J., Modo, M., Henry, L., & Gonzalez-Martinez, J. A. (2025). Structural connectivity changes in focal epilepsy: Beyond the epileptogenic zone. Epilepsia, 66(1), 226–239. 10.1111/EPI.18175

Behanova, A., Abdollahzadeh, A., Belevich, I., Jokitalo, E., Sierra, A., & Tohka, J. (2022). gACSON software for automated segmentation and morphology analyses of myelinated axons in 3D electron microscopy. Computer Methods and Programs in Biomedicine, 220, 106802. 10.1016/j.cmpb.2022.106802

Belevich, I., Joensuu, M., Kumar, D., Vihinen, H., & Jokitalo, E. (2016). Microscopy Image Browser: A Platform for Segmentation and Analysis of Multidimensional Datasets. PLOS Biology, 14(1), e1002340. 10.1371/JOURNAL.PBIO.1002340

Bernasconi, A., Cendes, F., Theodore, W. H., Gill, R. S., Koepp, M. J., Hogan, R. E., Jackson, G. D., Federico, P., Labate, A., Vaudano, A. E., Blümcke, I., Ryvlin, P., & Bernasconi, N. (2019). Recommendations for the use of structural magnetic resonance imaging in the care of patients with epilepsy: A consensus report from the International League Against Epilepsy Neuroimaging Task Force. Epilepsia, 60(6), 1054–1068. 10.1111/EPI.15612

Blumcke, I., Spreafico, R., Haaker, G., Coras, R., Kobow, K., Bien, C. G., Pfäfflin, M., Elger, C., Widman, G., Schramm, J., Becker, A., Braun, K. P., Leijten, F., Baayen, J. C., Aronica, E., Chassoux, F., Hamer, H., Stefan, H., Rössler, K., … Avanzini, G. (2017). Histopathological Findings in Brain Tissue Obtained during Epilepsy Surgery. New England Journal of Medicine, 377(17), 1648–1656. 10.1056/nejmoa1703784

Buchin, A., de Frates, R., Nandi, A., Mann, R., Chong, P., Ng, L., Miller, J., Hodge, R., Kalmbach, B., Bose, S., Rutishauser, U., McConoughey, S., Lein, E., Berg, J., Sorensen, S., Gwinn, R., Koch, C., Ting, J., & Anastassiou, C. A. (2022). Multi-modal characterization and simulation of human epileptic circuitry. Cell Reports, 41(13), 111873. 10.1016/j.celrep.2022.111873

Carmeli, C., Bonifazi, P., Robinson, H. P. C., & Small, M. (2013). Quantifying network properties in multi-electrode recordings: Spatiotemporal characterization and inter-trial variation of evoked gamma oscillations in mouse somatosensory cortex in vitro. Frontiers in Computational Neuroscience, 7(OCT), 59968. 10.3389/FNCOM.2013.00134/BIBTEX

Dastgheyb, R. M., Yoo, S.-W., & Haughey, N. J. (2020). MEAnalyzer – a Spike Train Analysis Tool for Multi Electrode Arrays. Neuroinformatics, 18(1), 163–179. 10.1007/s12021-019-09431-0

De Ciantis, A., Barba, C., Tassi, L., Cosottini, M., Tosetti, M., Costagli, M., Bramerio, M., Bartolini, E., Biagi, L., Cossu, M., Pelliccia, V., Symms, M. R., & Guerrini, R. (2016). 7T MRI in focal epilepsy with unrevealing conventional field strength imaging. Epilepsia, 57(3), 445–454. 10.1111/epi.13313

Dougalis, A., Abushik, P., Pelkonen, A., Giudice, L., Gómez-Budia, M., Novosolova, N., Välimäki, N.-N., Rezaie, M., Nurkhametova, D., Giniatullina, R., Shakirzyanova, A., Mali, A., Rauramaa, T., Stevens, B., Hiltunen, M., Leinonen, V., & Malm, T. (2025). Early amyloid spine response and impaired synaptic transmission of pyramidal neurons in human biopsies with Alzheimer’s Disease-related pathology. BioRxiv, 2025.01.07.630516. 10.1101/2025.01.07.630516

Dougalis, A. G., Matthews, G. A. C., Bishop, M. W., Brischoux, F., Kobayashi, K., & Ungless, M. A. (2012). Functional properties of dopamine neurons and co-expression of vasoactive intestinal polypeptide in the dorsal raphe nucleus and ventro-lateral periaqueductal grey. The European Journal of Neuroscience, 36(10), 3322. 10.1111/J.1460-9568.2012.08255.X

Fagerlund, I., Dougalis, A., Shakirzyanova, A., Gómez-Budia, M., Pelkonen, A., Konttinen, H., Ohtonen, S., Fazaludeen, M. F., Koskuvi, M., Kuusisto, J., Hernández, D., Pebay, A., Koistinaho, J., Rauramaa, T., Lehtonen, Š., Korhonen, P., & Malm, T. (2022). Microglia-like cells promote neuronal functions in cerebral organoids. Cells, 11(1), 124. 10.3390/CELLS11010124/S1

Gazestani, V., Kamath, T., Nadaf, N. M., Dougalis, A., Burris, S. J., Rooney, B., Junkkari, A., Vanderburg, C., Pelkonen, A., Gomez-Budia, M., Välimäki, N.-N., Rauramaa, T., Therrien, M., Koivisto, A. M., Tegtmeyer, M., Herukka, S.-K., Abdulraouf, A., Marsh, S. E., Hiltunen, M., … Macosko, E. Z. (2023). Early Alzheimer’s disease pathology in human cortex involves transient cell states. Cell, 186(20), 4438–4453.e23. 10.1016/j.cell.2023.08.005

Gorgolewski, K. J., Auer, T., Calhoun, V. D., Craddock, R. C., Das, S., Duff, E. P., Flandin, G., Ghosh, S. S., Glatard, T., Halchenko, Y. O., Handwerker, D. A., Hanke, M., Keator, D., Li, X., Michael, Z., Maumet, C., Nichols, B. N., Nichols, T. E., Pellman, J., … Poldrack, R. A. (2016). The brain imaging data structure, a format for organizing and describing outputs of neuroimaging experiments. Scientific Data, 3(1), 160044. 10.1038/sdata.2016.44

Hakkarainen, H., Sierra, A., Mangia, S., Garwood, M., Michaeli, S., Gröhn, O., & Liimatainen, T. (2016). MRI relaxation in the presence of fictitious fields correlates with myelin content in normal rat brain. Magnetic Resonance in Medicine, 75(1), 161–168. 10.1002/mrm.25590

Jehi, L. (2018). The epileptogenic zone: Concept and definition. Epilepsy Currents, 18(1), 12–16. 10.5698/1535-7597.18.1.12/ASSET/IMAGES/LARGE/10.5698_1535-7597.18.1.12-FIG3.JPEG

Kälviäinen, R., Hadj-Allal, Z., Kirjavainen, J., Roivainen, R., Linnankivi, T., Peltola, J., Eriksson, K., Lamusuo, S., Lähdesmäki, T., Annunen, J., Vieira, P., Tarkiainen, V., Jutila, L., Saarela, A., Kämppi, L., Metsähonkala, L., Gaily, E., Lähde, N., Antinmaa, J., … Sorjonen, P. (2025). Epilepsy care pathway: The Finnish model. Epilepsia Open, 10(1), 177–185. 10.1002/EPI4.13093

Kananen, J., Helakari, H., Korhonen, V., Huotari, N., Järvelä, M., Raitamaa, L., Raatikainen, V., Rajna, Z., Tuovinen, T., Nedergaard, M., Jacobs, J., LeVan, P., Ansakorpi, H., & Kiviniemi, V. (2020). Respiratory-related brain pulsations are increased in epilepsy—a two-centre functional MRI study. Brain Communications, 2(2), fcaa076. 10.1093/braincomms/fcaa076

Kobow, K., Ziemann, M., Kaipananickal, H., Khurana, I., Mühlebner, A., Feucht, M., Hainfellner, J. A., Czech, T., Aronica, E., Pieper, T., Holthausen, H., Kudernatsch, M., Hamer, H., Kasper, B. S., Rössler, K., Conti, V., Guerrini, R., Coras, R., Blümcke, I., … Kaspi, A. (2019). Genomic DNA methylation distinguishes subtypes of human focal cortical dysplasia. Epilepsia, 60(6), 1091–1103. 10.1111/epi.14934

Kumar, P., Lim, A., Hazirah, S. N., Chua, C. J. H., Ngoh, A., Poh, S. L., Yeo, T. H., Lim, J., Ling, S., Sutamam, N. B., Petretto, E., Low, D. C. Y., Zeng, L., Tan, E.-K., Arkachaisri, T., Yeo, J. G., Ginhoux, F., Chan, D., & Albani, S. (2022). Single-cell transcriptomics and surface epitope detection in human brain epileptic lesions identifies pro-inflammatory signaling. Nature Neuroscience, 25(7), 956–966. 10.1038/s41593-022-01095-5

Laufs, H. (2012). Functional imaging of seizures and epilepsy: evolution from zones to networks. Current Opinion in Neurology, 25(2), 194–200. 10.1097/WCO.0B013E3283515DB9

Lee, B. R., Dalley, R., Miller, J. A., Chartrand, T., Close, J., Mann, R., Mukora, A., Ng, L., Alfiler, L., Baker, K., Bertagnolli, D., Brouner, K., Casper, T., Csajbok, E., Donadio, N., Driessens, S. L. W., Egdorf, T., Enstrom, R., Galakhova, A. A., … Ting, J. T. (2023). Signature morphoelectric properties of diverse GABAergic interneurons in the human neocortex. Science, 382(6667). 10.1126/SCIENCE.ADF6484/SUPPL_FILE/SCIENCE.ADF6484_MDAR_REPRODUCIBILITY_CHECKLIST.PDF

Lehto, L. J., Albors, A. A., Sierra, A., Tolppanen, L., Eberly, L. E., Mangia, S., Nurmi, A., Michaeli, S., & Gröhn, O. (2017). Lysophosphatidyl Choline Induced Demyelination in Rat Probed by Relaxation along a Fictitious Field in High Rank Rotating Frame. Frontiers in Neuroscience, 11, 433. 10.3389/fnins.2017.00433

Luhmann, H. J., Dzhala, V. I., & Ben-Ari, Y. (2000). Generation and propagation of 4-AP-induced epileptiform activity in neonatal intact limbic structures in vitro. The European Journal of Neuroscience, 12(8), 2757–2768. 10.1046/j.1460-9568.2000.00156.x

Mohan, H., Verhoog, M. B., Doreswamy, K. K., Eyal, G., Aardse, R., Lodder, B. N., Goriounova, N. A., Asamoah, B., B. Brakspear, A. B. C., Groot, C., Van Der Sluis, S., Testa-Silva, G., Obermayer, J., Boudewijns, Z. S. R. M., Narayanan, R. T., Baayen, J. C., Segev, I., Mansvelder, H. D., & De Kock, C. P. J. (2015). Dendritic and Axonal Architecture of Individual Pyramidal Neurons across Layers of Adult Human Neocortex. Cerebral Cortex (New York, N.Y. : 1991), 25(12), 4839–4853. 10.1093/CERCOR/BHV188

Muhlhofer, W., Tan, Y. L., Mueller, S. G., & Knowlton, R. (2017). MRI-negative temporal lobe epilepsy—What do we know? Epilepsia, 58(5), 727–742. 10.1111/EPI.13699

Narvaez, O., Svenningsson, L., Yon, M., Sierra, A., & Topgaard, D. (2022). Massively Multidimensional Diffusion-Relaxation Correlation MRI. Frontiers in Physics, 9. 10.3389/fphy.2021.793966

Narvaez, O., Yon, M., Jiang, H., Bernin, D., Forssell-Aronsson, E., Sierra, A., & Topgaard, D. (2024). Nonparametric distributions of tensor-valued Lorentzian diffusion spectra for model-free data inversion in multidimensional diffusion MRI. Journal of Chemical Physics, 161(8), 84201. 10.1063/5.0213252/3309344

Pai, B., Tome-Garcia, J., Cheng, W. S., Nudelman, G., Beaumont, K. G., Ghatan, S., Panov, F., Caballero, E., Sarpong, K., Marcuse, L., Yoo, J., Jiang, Y., Schaefer, A., Akbarian, S., Sebra, R., Pinto, D., Zaslavsky, E., & Tsankova, N. M. (2022). High-resolution transcriptomics informs glial pathology in human temporal lobe epilepsy. Acta Neuropathologica Communications, 10(1), 149. 10.1186/s40478-022-01453-1

Pfisterer, U., Petukhov, V., Demharter, S., Meichsner, J., Thompson, J. J., Batiuk, M. Y., Asenjo-Martinez, A., Vasistha, N. A., Thakur, A., Mikkelsen, J., Adorjan, I., Pinborg, L. H., Pers, T. H., von Engelhardt, J., Kharchenko, P. V, & Khodosevich, K. (2020). Identification of epilepsy-associated neuronal subtypes and gene expression underlying epileptogenesis. Nature Communications, 11(1), 5038. 10.1038/s41467-020-18752-7

Rigas, P., Adamos, D. A., Sigalas, C., Tsakanikas, P., Laskaris, N. A., & Skaliora, I. (2015). Spontaneous Up states in vitro: a single-metric index of the functional maturation and regional differentiation of the cerebral cortex. Frontiers in Neural Circuits, 9, 59. 10.3389/fncir.2015.00059

Romigi, A., Bari, M., Placidi, F., Marciani, M. G., Malaponti, M., Torelli, F., Izzi, F., Prosperetti, C., Zannino, S., Corte, F., Chiaramonte, C., & Maccarrone, M. (2010). Cerebrospinal fluid levels of the endocannabinoid anandamide are reduced in patients with untreated newly diagnosed temporal lobe epilepsy. Epilepsia, 51(5), 768–772. 10.1111/j.1528-1167.2009.02334.x

Salo, R. A., Belevich, I., Jokitalo, E., Gröhn, O., & Sierra, A. (2021). Assessment of the structural complexity of diffusion MRI voxels using 3D electron microscopy in the rat brain. NeuroImage, 225, 117529. 10.1016/j.neuroimage.2020.117529

Sanders, M. W., Van der Wolf, I., Jansen, F. E., Aronica, E., Helmstaedter, C., Racz, A., Surges, R., Grote, A., Becker, A. J., Rheims, S., Catenoix, H., Duncan, J. S., De Tisi, J., Jacques, T. S., Cross, J. H., Kalviainen, R., Rauramaa, T., Chassoux, F., Devaux, B. C., … Braun, K. P. (2024). Outcome of Epilepsy Surgery in MRI-Negative Patients Without Histopathologic Abnormalities in the Resected Tissue. Neurology, 102(4). 10.1212/WNL.0000000000208007/SUPPL_FILE/SUPPLEMENTARY_TAB LE1.PDF

Peddie, C.J., Genoud, C., Kreshuk, A., Meechan, K., Micheva, K.D., Narayan, K., Pape, C., Parton, R.G., Schieber, N.L., Schwab, Y., Titze, B., Verkade, P., Aubrey, A., & Collinson, L.M.. (2022). Volume electron microscopy. Nat Rev Methods Primers. 7;2:51. 10.1038/s43586-022-00131-9.

Scoyni, F., Giudice, L., Väänänen, M.-A., Downes, N., Korhonen, P., Choo, X. Y., Välimäki, N.-N., Mäkinen, P., Korvenlaita, N., Rozemuller, A. J., de Vries, H. E., Polo, J., Turunen, T. A., Ylä-Herttuala, S., Hansen, T. B., Grubman, A., Kaikkonen, M. U., & Malm, T. (2024). Alzheimer’s disease-induced phagocytic microglia express a specific profile of coding and non-coding RNAs. Alzheimer’s & Dementia, 20(2), 954–974. 10.1002/alz.13502

Spitzer, H., Ripart, M., Whitaker, K., D’Arco, F., Mankad, K., Chen, A. A., Napolitano, A., De Palma, L., De Benedictis, A., Foldes, S., Humphreys, Z., Zhang, K., Hu, W., Mo, J., Likeman, M., Davies, S., Güttler, C., Lenge, M., Cohen, N. T., … Wagstyl, K. (2022). Interpretable surface-based detection of focal cortical dysplasias: a Multi-centre Epilepsy Lesion Detection study. Brain, 145(11), 3859–3871. 10.1093/BRAIN/AWAC224

Straehle, J., Ravi, V. M., Heiland, D. H., Galanis, C., Lenz, M., Zhang, J., Neidert, N. N., El Rahal, A., Vasilikos, I., Kellmeyer, P., Scheiwe, C., Klingler, J. H., Fung, C., Vlachos, A., Beck, J., & Schnell, O. (2023). Technical report: surgical preparation of human brain tissue for clinical and basic research. Acta Neurochirurgica, 165(6), 1461–1471. 10.1007/s00701-023-05611-9

Stuart, T., Butler, A., Hoffman, P., Hafemeister, C., Papalexi, E., Mauck, W. M., Hao, Y., Stoeckius, M., Smibert, P., & Satija, R. (2019). Comprehensive Integration of Single-Cell Data. Cell, 177(7), 1888–1902.e21. 10.1016/j.cell.2019.05.031

Tóth, K., Hofer, K. T., Kandrács, Á., Entz, L., Bagó, A., Erőss, L., Jordán, Z., Nagy, G., Sólyom, A., Fabó, D., Ulbert, I., & Wittner, L. (2018). Hyperexcitability of the network contributes to synchronization processes in the human epileptic neocortex. The Journal of Physiology, 596(2), 317–342. 10.1113/JP275413

van Lanen, R. H. G. J., Colon, A. J., Wiggins, C. J., Hoeberigs, M. C., Hoogland, G., Roebroeck, A., Ivanov, D., Poser, B. A., Rouhl, R. P. W., Hofman, P. A. M., Jansen, J. F. A., Backes, W., Rijkers, K., & Schijns, O. E. M. G. (2021). Ultra-high field magnetic resonance imaging in human epilepsy: A systematic review. NeuroImage: Clinical, 30, 102602. 10.1016/J.NICL.2021.102602

Wierda, K., Nyitrai, H., Lejeune, A., Vlaeminck, I., Leysen, E., Theys, T., de Wit, J., Vanderhaeghen, P., & Libé-Philippot, B. (2024). Protocol to process fresh human cerebral cortex biopsies for patch-clamp recording and immunostaining. STAR Protocols, 5(4). 10.1016/J.XPRO.2024.103313

Wilkinson, M. D., Dumontier, M., Aalbersberg, Ij. J., Appleton, G., Axton, M., Baak, A., Blomberg, N., Boiten, J.-W., da Silva Santos, L. B., Bourne, P. E., Bouwman, J., Brookes, A. J., Clark, T., Crosas, M., Dillo, I., Dumon, O., Edmunds, S., Evelo, C. T., Finkers, R., … Mons, B. (2016). The FAIR Guiding Principles for scientific data management and stewardship. Scientific Data, 3(1), 160018. 10.1038/sdata.2016.18

Wirrell, E. C., Nabbout, R., Scheffer, I. E., Alsaadi, T., Bogacz, A., French, J. A., Hirsch, E., Jain, S., Kaneko, S., Riney, K., Samia, P., Snead, O. C., Somerville, E., Specchio, N., Trinka, E., Zuberi, S. M., Balestrini, S., Wiebe, S., Cross, J. H., … Tinuper, P. (2022). Methodology for classification and definition of epilepsy syndromes with list of syndromes: Report of the ILAE Task Force on Nosology and Definitions. Epilepsia, 63(6), 1333–1348. 10.1111/EPI.17237

Yang, M., Zhang, Y., Zhang, T., Zhou, H., Ren, J., Cao, X., Zhou, D., & Yang, T. (2024). Postoperative seizure outcomes and antiseizure medication utilization based on histopathological diagnosis: A retrospective cohort study. Epilepsy & Behavior, 161, 110056. 10.1016/J.YEBEH.2024.110056

